# Subchronic Toxicity Assessment of Perfluoroalkyl Substances by Mixed Exposure of Nine Substances at Daily Intake Relevant Concentration

**DOI:** 10.1101/2023.05.17.541082

**Authors:** Kazuki Takeda, Taki Saito, Sakura Sasaki, Akifumi Eguchi, Makoto Sugiyama, Kio Suzuki, Saeka Eto, Ryo Kamata

**Affiliations:** Laboratory of Toxicology, School of Veterinary Medicine, Kitasato University, E23-35-1, Towada, Aomori 034-0021, Japan; Department of Computer Science, Tokyo Institute of Technology, 4259-J3-1818, Nagatsuta-cho, Midori-ku, Yokohama-shi, Kanagawa 226-0026, Japan; Laboratory of Toxicology, Department of Environmental Veterinary Sciences, Faculty of Veterinary Medicine, Hokkaido University, N18 W9, Kita-ku, Sapporo 060-0818, Japan; Department of Sustainable Health Science, Center for Preventive Medical Sciences, Chiba University, Chiba, Japan; Laboratory of Anatomy, School of Veterinary Medicine, Kitasato University, Aomori, Japan

## Abstract

Per- and poly-fluoroalkyl substances (PFAS) have been utilized extensively for various applications owing to their distinctive chemical properties. They exhibit high persistence in the environment and accumulate within the human body, necessitating toxicity assessments. However, the consequences of prolonged, low-level exposure to PFAS and concurrent exposure to multiple PFAS have not been explored. In this study, male C57BL/6J mice (aged 8 weeks) were exposed to a composite of nine PFAS, which include long-chain PFAS (e.g. perfluorooctanoic acid and perfluorooctanesulfonic acid) and short-chain PFAS (e.g. perfluorobutanoic acid and perfluorobutanesulfonic acid) at concentrations equivalent to the estimated daily human intake in the composition reported (1 µg/L [sum of the nine compounds], the maximum reported exposure concentration) via drinking water. Histological examination revealed vacuolization of hepatocytes and irregular arrangement of hepatocyte cords, suggesting that exposure to low levels of the PFAS mixture causes morphological changes in liver tissues. Transcriptome analysis revealed that PFAS exposure mainly altered a group of genes related to metabolism and chemical carcinogenesis. Machine learning analysis of the liver metabolome showed a typical concentration-independent alteration upon PFAS exposure, and in addition to known substances such as glutathione, a compound with unknown biological function; 2,5-dihydro-2,4-dimethyloxazole was found. This study demonstrates that daily exposure to PFAS leads to morphological changes in liver tissues and alters the expression of metabolism-and cancer-related genes as well as phospholipid metabolism. Future studies are required to evaluate the chronic toxicity of prolonged, low-level exposure to PFAS mixtures and to investigate the health effects of PFAS.

## 1. Introduction

Per-and poly-fluoroalkyl substances (PFAS) include thousands of chemical compounds containing the perfluoroalkyl moiety C_n_F_2n+1_– (Buck et al., 2011). It is estimated that more than 1,000,000 tons of PFAS including hydrofluorocarbons are produced annually worldwide (Evich et al., 2022). Their exclusive chemical properties, such as resistance to heat and chemicals, water and oil repellency, emulsifying characteristics, and light absorption, accredit a broad spectrum of applications. However, PFAS are exceedingly persistent in the environment and bioaccumulate easily in the body; they are therefore referred to as “Forever Chemicals,” raising concerns about their health implications (David Kempisty and Racz, 2021). Consequently, PFAS contamination has gained global attention, with McDonald’s Corporation pledging to eradicate PFAS from food packaging within the next 5 years and Amazon Corporation announcing in December of the same year a prohibition on using PFAS in food containers (Stohler, 2020).

Among the different classes of PFAS, perfluorooctanoic acid (PFOA) and perfluorooctanesulfonic acid (PFOS) are of significant concern owing to their high bioaccumulation potential. The geometric mean human serum elimination half-life of PFOS is 4.8 years (95% confidence interval [CI], 4.0–5.8) and that of PFOA is 3.5 years (95% CI, 3.0–4.1) (Olsen et al., 2007). The production and use of these substances are regulated internationally. PFOS was included in Annex B of the Stockholm Convention on Persistent Organic Pollutants in 2009, and its production, use, import, and export are generally restricted. PFOA and its related compounds were listed in Annex A in 2019 and perfluorohexane sulfonic acid (PFHxS) was listed in Annex A in 2022 (Stockholm Convention, 2022). Safety thresholds have been set for major PFAS. The European Food Safety Authority (EFSA) has set the derived no-effect level (DNEL) for PFOA and PFOS at 1.5 µg/kg/day and 0.15 µg/kg/day, respectively (Alexander et al., 2008). In contrast, the U.S. Environmental Protection Agency (EPA) has set the DNEL for PFOA and PFOS at 0.1 µg/kg/day and 0.03 µg/kg/day, respectively. These limits have been regulated more tightly in recent years, and EFSA set a new total weekly tolerable intake of 4.4 ng/kg BW/week for PFOA, PFOS, perfluorononanoic acid (PFNA), and PFHxS in 2020 (Schrenk et al., 2020). Furthermore, the EPA’s lifetime health advisory level, the level at which no health effects are expected after 70 years of drinking 2 L of water per day, was set at 70 ng/L for PFOA and PFOS combined in 2016. However, it was lowered to 0.024 ng/L in June 2022 (US EPA, 2022). In Japan, the Ministry of Health, Labour and Welfare (MHLW) listed PFOS and PFOA as water quality management targets in April 2020 and set a preliminary target of 50 ng/L for both compounds combined (Japanese Ministry of the Environment, 2020). However, regulatory levels for PFOA and PFOS vary widely among countries and organizations. The MHLW has set the standard levels for PFOS and PFOA as provisional guideline levels; therefore, no conclusion based on sufficient scientific evidence has been reached as to whether these limits are appropriate.

Although the complete extent of the adverse outcome pathway (AOP) of PFAS has not been comprehensively understood, the detrimental effects of PFAS are known to be characterized by hepatotoxicity, lipid metabolism disruption, hypothyroidism, immunosuppression, reproductive toxicity, and carcinogenicity (Fenton et al., 2021). A previous study reported that one of the mechanisms of PFAS toxicity involves the activation of nuclear receptors, such as peroxisome proliferator-activated receptor (PPAR) α, PPARγ, PPARδ, constitutive androstane receptor, and pregnane X receptor (PXR) (Bjork et al., 2011). Additionally, activation of PXR and other receptors has been found to contribute to disease pathogenesis (Rosen et al., 2017). The liver, where these nuclear receptors are active, is the primary target for toxicity. In mice, dose-dependent liver weight increase, hepatocyte hypertrophy with vacuole formation, and increased (or loss of) peroxisome proliferation were observed at high body burdens of long-chain PFAS (Blake et al., 2020). Studies using cultured cells have indicated a link between PFAS and cancer; for example, PFOA, PFOS, and PFHxS have been reported to induce cell proliferation and malignant transformation in human mammary epithelial cells (Pierozan et al., 2022, 2020). Although short-chain PFAS such as perfluorobutanoic acid (PFBA) and perfluorobutanesulfonic acid (PFBS) have been used as alternatives to long-chain PFAS, drinking water exposure studies of PFBS in mice have found that it alters the liver and intestinal metabolome (Chen et al., 2023, 2022). However, most of these prior studies were conducted based on exposure to relatively high concentrations of single compounds.

The long half-lives of PFAS in the environment and *in vivo* suggest that people are exposed to a wide range of PFAS at lower concentrations and in more diverse forms over prolonged periods. There is relatively limited evidence from animal studies that replicate such real-world exposure conditions. Exposure to a PFAS mixture that mimicked an environment contaminated with PFAS in ski wax altered dopamine levels in mice (Grønnestad et al., 2021). Moreover, exposure to PFOS at 2 ng/g resulted in stunted embryos in bovine cumulus oocytes (Hallberg et al., 2021). However, little is known about the toxic effects of PFAS at concentrations that humans may ingest daily.

Mixed exposure is expected to occur in actual exposure situations as multiple PFAS remain in the environment. As the sources of exposure are diverse, including drinking water, food, and dust, it is challenging to estimate the exact amount of exposure. Pérez et al. (2014) assessed PFAS concentrations in food in different regions worldwide and estimated the daily human intake of PFAS by region. The concentration range was broad, ranging 30–100 ng/kg/day, with the highest estimate being that children around the Mediterranean Sea may have ingested nearly 100 ng/kg/day of PFAS. Therefore, herein, we set 100 ng/kg/day as the maximum anticipated daily intake in humans and evaluated the toxic effects of nine linear PFAS, including long-chain PFAS (25 ng/kg/day of PFOA and 15 ng/kg/day of PFOS) and short-chain PFAS (20 ng/kg/day of PFBA and 6 ng/kg/day of PFBS), in mice via drinking water exposure for 5 weeks. We hypothesized that toxic effects could occur in mixed exposures even if each individual substance is at a low concentration and includes a short-chain PFAS. Additionally, histological examination was performed to observe subtle toxic effects that cannot be confirmed at the organism level. Liver, where significant histological changes occurred, was subjected to multi-omics (transcriptome and metabolome) analysis to elucidate the mechanism of toxicity. This study provides scientific insight into the toxic effects of low-concentration mixtures of PFAS to which humans can actually be exposed.

## 2. Material and Methods

### 2.1. Materials

Nine major linear-chain PFAS were tested in this study—PFOA, PFBA, PFOS, PFPeA, PFNA, PFHxA, PFBS, PFHpA, and PFHxS—and their exposure concentrations and compositions were in accordance with a rationale paper mentioned above (Pérez et al., 2014). The composition of the compounds is listed in Table 1. PFOA, PFOS, and PFHxS were purchased from Sigma-Aldrich (St. Louis, MO), whereas others were obtained from Tokyo Chemical Industry Co., Ltd. (Tokyo, Japan).

**Table 1.**
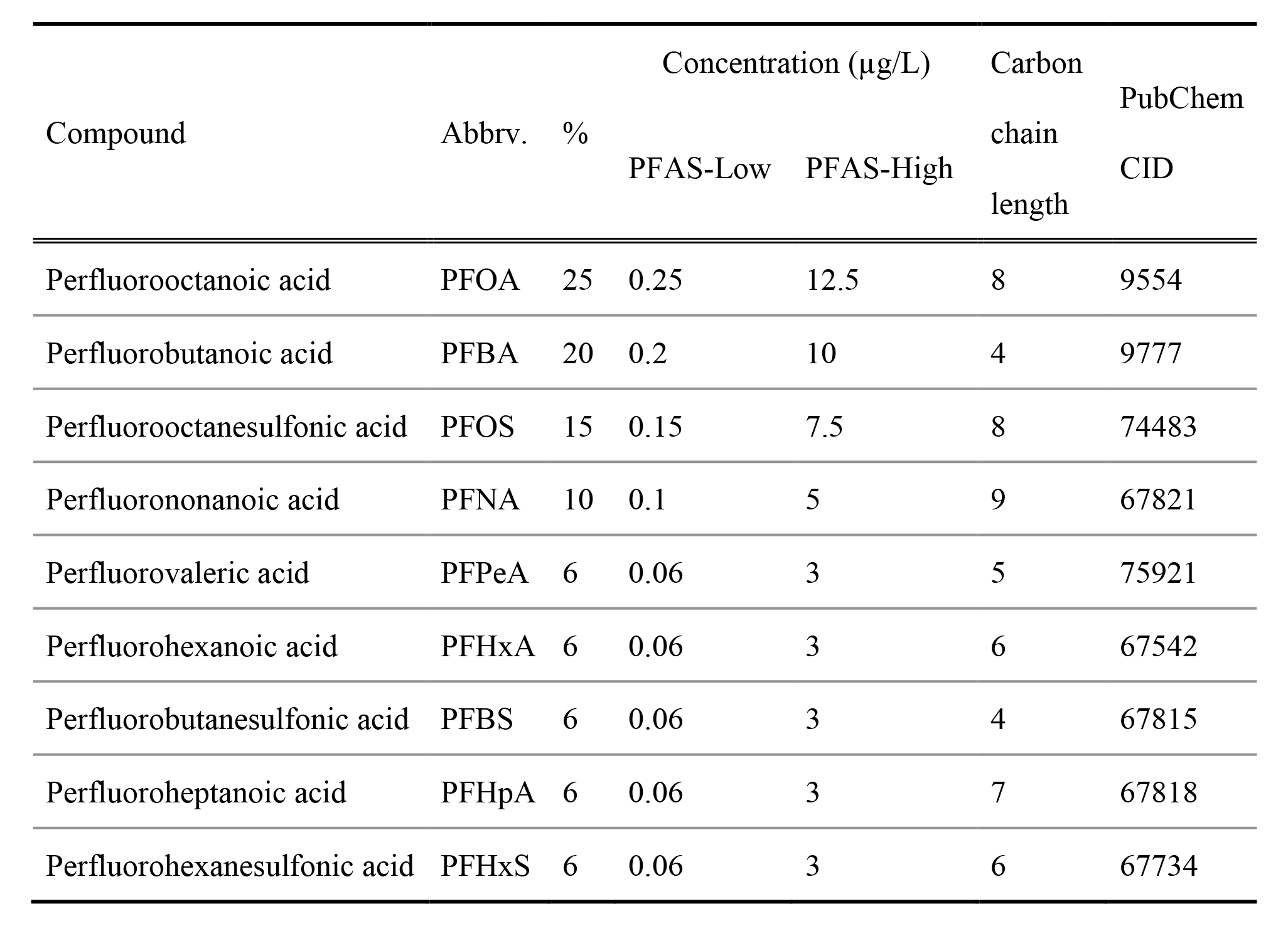
Composition of the nine poly-fluoroalkyl substances (PFAS) used in the study.

| Compound | Abbrev. | % | Concentration (µg/L) |  | Carbon chain length | PubChem CID |
| --- | --- | --- | --- | --- | --- | --- |
|  |  |  | PFAS-Low | PFAS-High |  |  |
| Perfluorooctanoic acid | PFOA | 25 | 0.25 | 12.5 | 8 | 9554 |
| Perfluorobutanoic acid | PFBA | 20 | 0.2 | 10 | 4 | 9777 |
| Perfluorooctanesulfonic acid | PFOS | 15 | 0.15 | 7.5 | 8 | 74483 |
| Perfluorononanoic acid | PFNA | 10 | 0.1 | 5 | 9 | 67821 |
| Perfluorovaleric acid | PFPeA | 6 | 0.06 | 3 | 5 | 75921 |
| Perfluorohexanoic acid | PFHxA | 6 | 0.06 | 3 | 6 | 67542 |
| Perfluorobutanesulfonic acid | PFBS | 6 | 0.06 | 3 | 4 | 67815 |
| Perfluoroheptanoic acid | PFHpA | 6 | 0.06 | 3 | 7 | 67818 |
| Perfluorohexanesulfonic acid | PFHxS | 6 | 0.06 | 3 | 6 | 67734 |

### 2.2. Animal experiment

Seven-week-old male C57BL/6J mice were purchased from CLEA Japan, Inc. (Tokyo, Japan). They were housed under a 12/12 hour light/dark cycle at 20°C–23°C. Food (CE-2; CLEA) and water gel (HydroGel, ClearH2O, Westbrook, ME) were provided *ad libitum* and the mice were not fasted before or during the experiments. Health observations were made thrice per week, and cages, food, and HydroGel were changed once per week. Following 1 week of adaptation, the mice were randomly divided into three groups, with five individuals in each group.

One group was treated with clean HydroGel (control group), while the other two were treated with HydroGel containing nine mixtures of PFAS at total concentrations of 1 µg/L and 50 µg/L (PFAS-Low and PFAS-High groups, respectively). The average daily intake of each mouse in the 1 µg/L of PFAS mix-and 50 µg/L of PFAS mix-treated groups was estimated at approximately 100 ng/kg/day and 5,000 ng/kg/day, respectively, since the average body weight at the beginning of the experiment was 30 g and the average daily drinking volume was 3 mL. After the 5-week exposure period, mice were euthanized by inhalation of an excess amount of isoflurane and laparotomized. The organs were harvested after complete blood sampling from the posterior vena cava. All animal care and experimental procedures were approved by the Animal Care Committee of Kitasato University School of Veterinary Medicine (Approval No. 21-012) and were conducted in accordance with this committee and national regulations.

### 2.3. Tissue histology

Four organs (liver, spleen, kidney, and heart) were immersed and fixed in 10% neutral formalin phosphate buffer solution for at least 1 month and then embedded in Pathoprep568 (Fujifilm Wako Pure Chemical Corporation, Tokyo, Japan), following routine methods. The paraffin blocks were cut into thin sections (4 µm) and stained with hematoxylin and eosin for morphological observation or with periodic acid-Schiff (PAS) with/without amylase digestion for glycogen detection in liver sections according to the method of Kovac et al. (2015).

### 2.4. Liver transcriptome analysis

A small piece of liver immersed in RNAlater™ solution (Sigma-Aldrich) was used for RNA extraction. RNA extraction was performed using TRI reagent (Cosmo Bio Co., Ltd., Tokyo, Japan) and the Monarch Total RNA Miniprep Kit (New England BioLabs Inc., Ipswich, MA) according to the manufacturer’s instructions. The concentration of RNA in the extract was quantified by measuring the absorbance at 260 nm using the BioSpectrometer kinetic (Eppendorf, Hamburg, Germany). Subsequent RNA sequence analysis (RNA-seq) of the purified RNA was performed by GenScript Biotech Corp. (Piscataway, NJ). Briefly, RNA was randomly fragmented and reverse transcribed into cDNA, and adapter sequences were attached to both ends of the fragments. The fragments were amplified using polymerase chain reaction (PCR), and those with a size of 200–400 base pairs were selected and sequenced using the NovaSeq 6000 system (Illumina Inc., San Diego, CA). Detected changes in gene expression levels were analyzed using the Kyoto Encyclopedia of Genes and Genomes (KEGG) pathway (Kanehisa et al., 2023).

### 2.5. Real-time quantitative PCR

Gene expression levels of eight genes representing pathways that varied in the transcriptome analysis were confirmed by quantitative PCR (qPCR) according to our previous report (Kamata et al., 2022). Briefly, reverse transcription from mRNA was performed using the LunaScript® RT SuperMix Kit (New England BioLabs) following standard protocols. The cDNA was amplified using Luna® Universal qPCR Master Mix and corresponding primers for RT-qPCR. The primers were designed using the National Center for Biotechnology Information Primer designing tools. The primer information is shown in Table S1. The PCR conditions were as follows: initial denaturation, 95°C for 60 s; 40 cycles of denaturation, 95°C for 15 s; and annealing and extension, 60°C for 30 s. Real-time PCR was performed for each sample using the StepOnePlus™ Real-Time PCR System (Thermo Fisher Scientific). β-actin (*Actb*) was used as the internal control, and the fold change was calculated using the 2^−ΔΔCt^ method.

### 2.6. Metabolome analysis of liver

Metabolome analysis was performed with liver samples following a previous report with slight modifications (Saigusa et al., 2016). Briefly, 400 μL of methanol containing internal standards (25 nM N,N-diethyl-2-phenylacetamide and d-camphor-10-sulfonic acid) and 400 μL of ultrapure water were added to 100 mg of liver sample and homogenized using BioMasher® II equipped with PowerMasher® II (Nippi, Incorporated, Tokyo, Japan) to extract the metabolome. The homogenates were centrifuged at 14,000 × rpm for 5 min after adding 100 μL of methanol containing internal standards (100 nM of N,N-diethyl-2-phenylacetamide and d-camphor-10-sulfonic acid). After centrifuging, the supernatant was transferred to Amicon® Ultra-0.5 3 kDa filter columns (Merck Millipore) and centrifuged at 14,000 × rpm for 1 hour. The filtrates were transferred to glass vials for ultra-performance liquid chromatography quantitative time-of-flight mass spectrometry (UPLC/QTOF-MS) analysis. This analysis was performed using an ExionLC AD UPLC system interfaced with an X500R QToF-MS system (SCIEX, Framingham, MA) with electrospray ionization (ESI) operated in positive and negative ion modes. The LC conditions and MS parameters employed were in accordance with the previous report (Saigusa et al., 2016). A serum quality control (QC) sample for metabolome analysis was prepared by pooling and mixing the same volume of all serum samples. The QC and blank samples (ultrapure water + internal standard) were injected at an interval of 6–7 sample injections to identify the sample carryover and check for stability during the entire analytical sequence.

In this study, metabolomes were detected in at least 50% of the analyzed samples. The coefficient of variation values of 30% of metabolomes and annotation level 2 proposed by Schymanski et al. (2014) were used for data analysis. Peak heights were normalized by peak heights of internal standards and locally weighted least-square regression (locally estimated smoothing function) and cubic spline with QC samples. The metabolome data was analyzed using Mass Spectrometry-Data Independent AnaLysis (MS-DIAL) software version 4.90 (Tsugawa et al., 2015) and R statistical environment Ver 4. 2. The mass spectra were searched against a RIKEN library, MS-bank North America, NIST20 tandem mass spectrometry library, and human metabolome database (Wishart et al., 2018). From candidates with the total scores based on the isotope ratio and accurate mass MS/MS similarity calculated, the annotation was made for the candidate with the highest score, and the identification cutoff score was set at 80.

### 2.7. Molecular docking

One of the possible mechanisms of PFAS toxicity is binding to the nuclear receptors PPARα, PPARγ, and PPARδ (Bjork et al., 2011; Fenton et al., 2021). Molecular docking simulations were performed to evaluate the binding affinity between the nine PFAS and the three PPARs. The 3D molecular structures of the small molecules and each known PPAR ligand were obtained from PubChem. Their PubChem CIDs are listed in Table 1. They were further preprocessed by adding partial atomic charges for their structures determined by molecular mechanics minimization calculated using the Gasteiger method (Gasteiger and Marsili, 1978). The 3D structures of murine PPARα, PPARδ, and PPARγ were constructed using the protein 3D structure modeling algorithm Alphafold2 (Jumper et al., 2021). The protein structure was preprocessed by adding hydrogen atoms and energy minimization using the CHARMM force field (Brooks et al., 1983). The binding pocket of each PPAR was determined using the coordinates of the pocket with the largest volume identified by GHECOM (Grid-based HECOMi finder) (Kawabata, 2010). Molecular docking was performed using AutoDock Vina to determine the best docking poses and docking scores (Trott and Olson, 2010). The search area was a 25 Å × 25 Å × 25 Å grid box, and each calculation was performed 5000 times. The 3D structures obtained above were converted to 2D structures using LIGPLOT+ v.2.2.5 to compare the respective amino acid binding modes (Laskowski and Swindells, 2011).

### 2.8. Biomarkers of liver injury

Liver injury and lipid metabolism biomarkers were measured using plasma samples collected after euthanasia at the end of PFAS exposure. Alkaline phosphatase (ALP) and cholesterol levels were measured colorimetrically using LabAssay™ (FUJIFILM Wako Pure Chemical Corporation, Osaka, Japan) according to standard protocols. Aspartate aminotransferase (AST) level was measured colorimetrically using a method established by Sigma-Aldrich (MAK055).

### 2.9. Data analysis

Statistical analysis was performed using JMP Pro 16 (SAS Institute, Cary, NC, USA). Tukey’s HSD test was used to compare measurements between all groups; a *p*-value of <0.05 was considered statistically significant. For omics analysis, the false discovery rate (FDR) was applied to adjust *p*-values. R packages in R v.4.2 were used to analyze the results of the omics analysis. MetaboAnalyst R was used for enrichment analysis of the metabolome (Xia et al., 2009). To characterize the metabolomics after PFAS exposure, Random Forests, a machine learning classification method, was utilized. PFAS-Low and PFAS-High were grouped together as PFAS-exposed groups, and a binary classification model was constructed to discriminate between the control and PFAS-exposed groups based on metabolomics. The implementation was based on the scikit-learn decision tree library with 5-fold cross-validation. n_estimator was set to 1000 and max_depth was set to 3.

## 3. Results

### 3.1. Histological examination

Histological examination was performed on four organs (liver, spleen, kidney, and heart) to detect histological changes upon PFAS exposure. The most remarkable changes were observed in the liver, with an irregular alignment of the hepatic lamina in all PFAS-exposed individuals. The cytoplasm was vacuolated, granular, and pale (Fig. 1A). Other symptoms like inflammation or apoptosis were not observed in all samples. Glycogen degeneration is one of the causes of such granular degeneration. To confirm whether the granular degeneration was caused by glycogen, a PAS reaction with amylase digestion was performed. The results showed that before the amylase digestion test, the control group was diffusely positive for the PAS reaction and that the PFAS-exposed group had a decreased degree of reaction; however, after the amylase digestion test, all groups were negative for the PAS reaction (Fig. 1B). No distinct histological changes were observed in the spleen, kidney, and heart (Fig. S1). There were also no significant changes in body weight, clinical symptoms before euthanasia, or gross organ abnormalities.

**Fig. 1.**
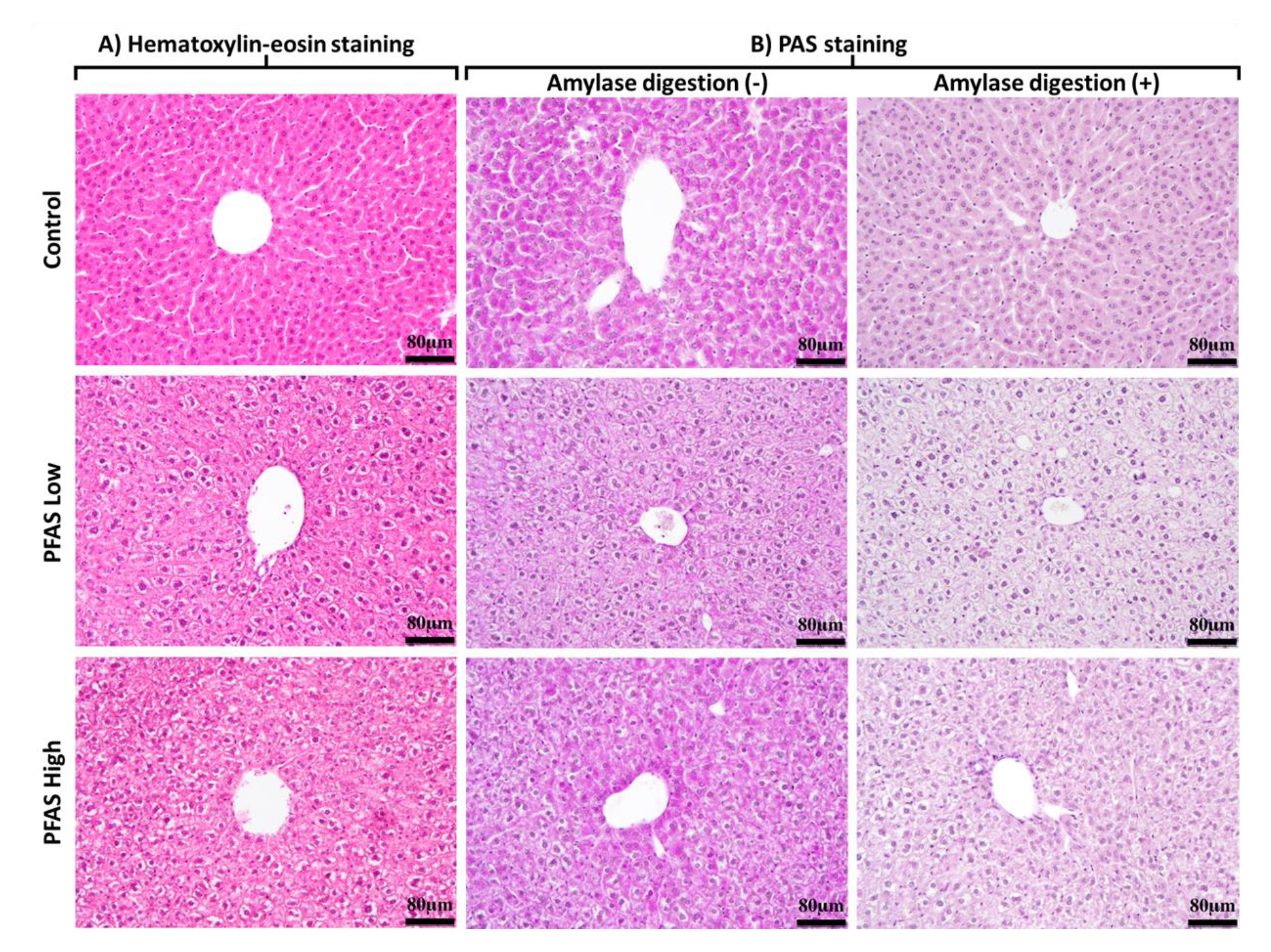
A) Histology of the liver determined by hematoxylin and eosin (HE) staining (magnification ×400) of samples in the control, poly-fluoroalkyl substances (PFAS)-Low (100 ng/kg/day), and PFAS-High (5,000 ng/kg/day) groups. Diffuse vacuolation and hypertrophy of hepatocytes with eosinophilic granules were observed in all mice in the PFAS-Low and -High groups. B) Representative images of livers stained with periodic acid-Schiff (PAS) stain with/without amylase digestion (magnification ×400).

### 3.2. Liver transcriptome analysis

Subsequent experiments focused on the liver, where significant histological changes were observed after PFAS exposure. Fig. 2A shows the number of genes with more than 2-fold changes in expression between each PFAS-exposed group detected by the RNA-seq. Clustering of these variations by Euclidean distance showed a similar pattern of gene variation in the PFAS-Low and PFAS-High groups compared with the control group (Fig. 2B). The fluctuating gene groups were classified by function based on KEGG pathways, and enrichment analysis was performed to show the number of significantly fluctuating genes (Fig. 2C and D). Compared with the control group, a total of 70 metabolism-related and 30 cancer-related genes were significantly altered in the PFAS-Low group and approximately 50 metabolism-related and 20 cancer-related genes were significantly altered in the PFAS-High group, with metabolism-related genes accounting for the highest number of alterations in the PFAS-Low and PFAS-High groups, followed by cancer-related genes.

**Fig. 2.**
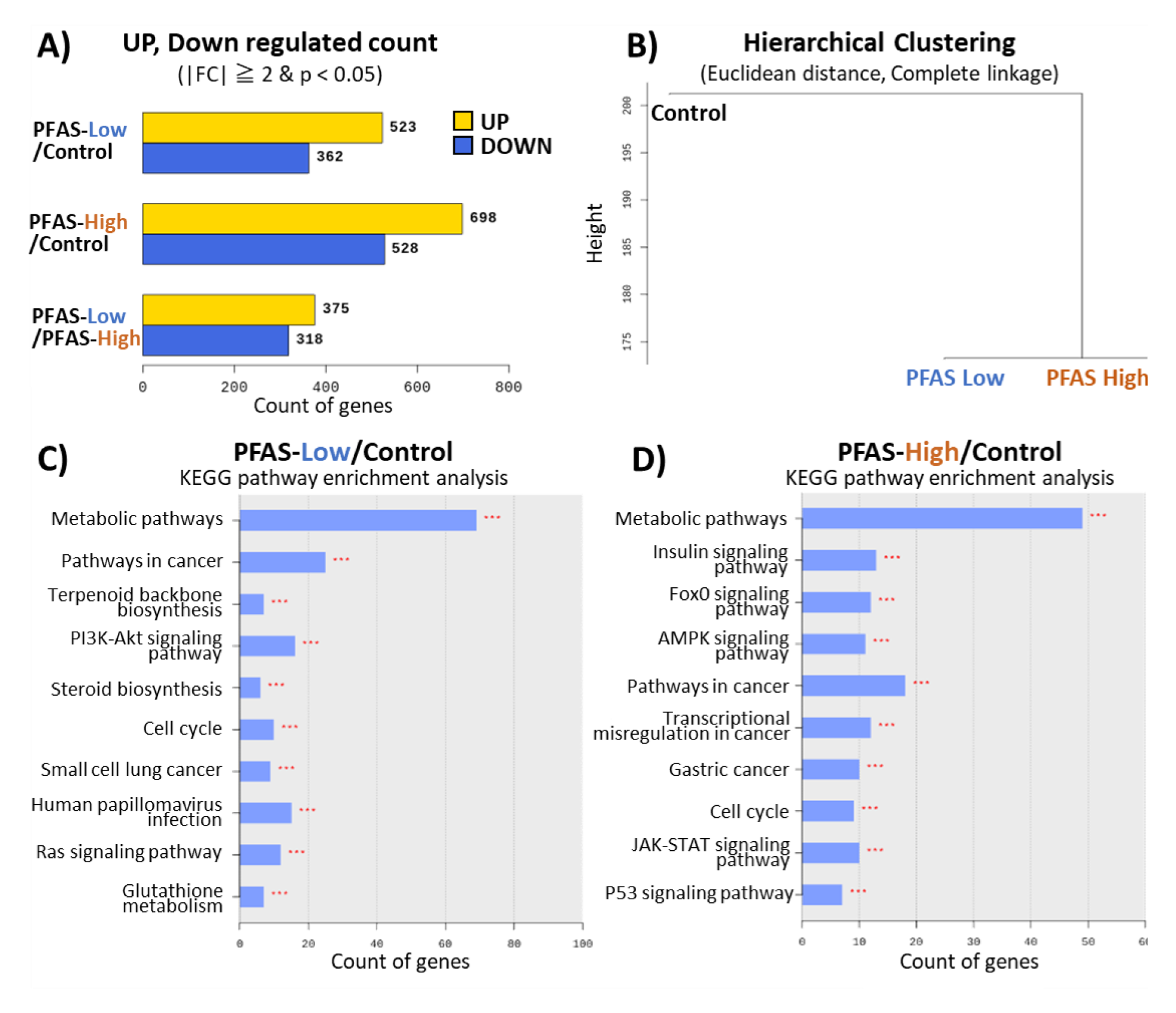
Transcriptome analysis of the liver. A) Shows the number of upregulated and downregulated genes based on the comparison pair’s fold change (FC). B) The high expression similarities were grouped together using each sample’s normalized value. (Distance metric = Euclidean distance, Linkage method = Complete Linkage). C) Kyoto Encyclopedia of Genes and Genomes (KEGG) enrichment analysis between the PFAS-Low and control groups. D) KEGG enrichment analysis between the PFAS-High and control groups.

### 3.3. qPCR

Based on enrichment analysis of the transcriptome, chemical carcinogenesis, cholesterol metabolism, and fatty acid metabolism were selected as pathways with several genes whose expression levels were significantly different in the liver due to PFAS exposure. Six representative target genes of each pathway (*Chrna4*, *Ccnd1*, *Pcsk9*, *Fasn*, *Cdc6*, and *Myc*) were quantified and validated in all individuals (Fig. 3). Compared with the control group, the most upregulated gene was *Chrna4*, with a 6-fold and 10-fold increase in expression in the PFAS-Low and PFAS-High groups, respectively (Tukey’s HSD test). Significant increases were observed for *Ccnd1*, a cell cycle regulator contributing to tumorigenesis, and *Fasn*, a fatty acid synthase promoting obesity and tumorigenesis in PFAS-exposed mice. *Pcsk9*, an aggravating factor for blood cholesterol levels, was only upregulated in the PFAS-Low group. Conversely, *Cdc6*, an oncogenic gene acting as a regulator of DNA replication, was significantly upregulated in only the PFAS-High group. Expression levels of *Myc* (c-Myc), an oncogenic gene, did not differ significantly between the control and PFAS-exposed groups.

**Fig. 3.**
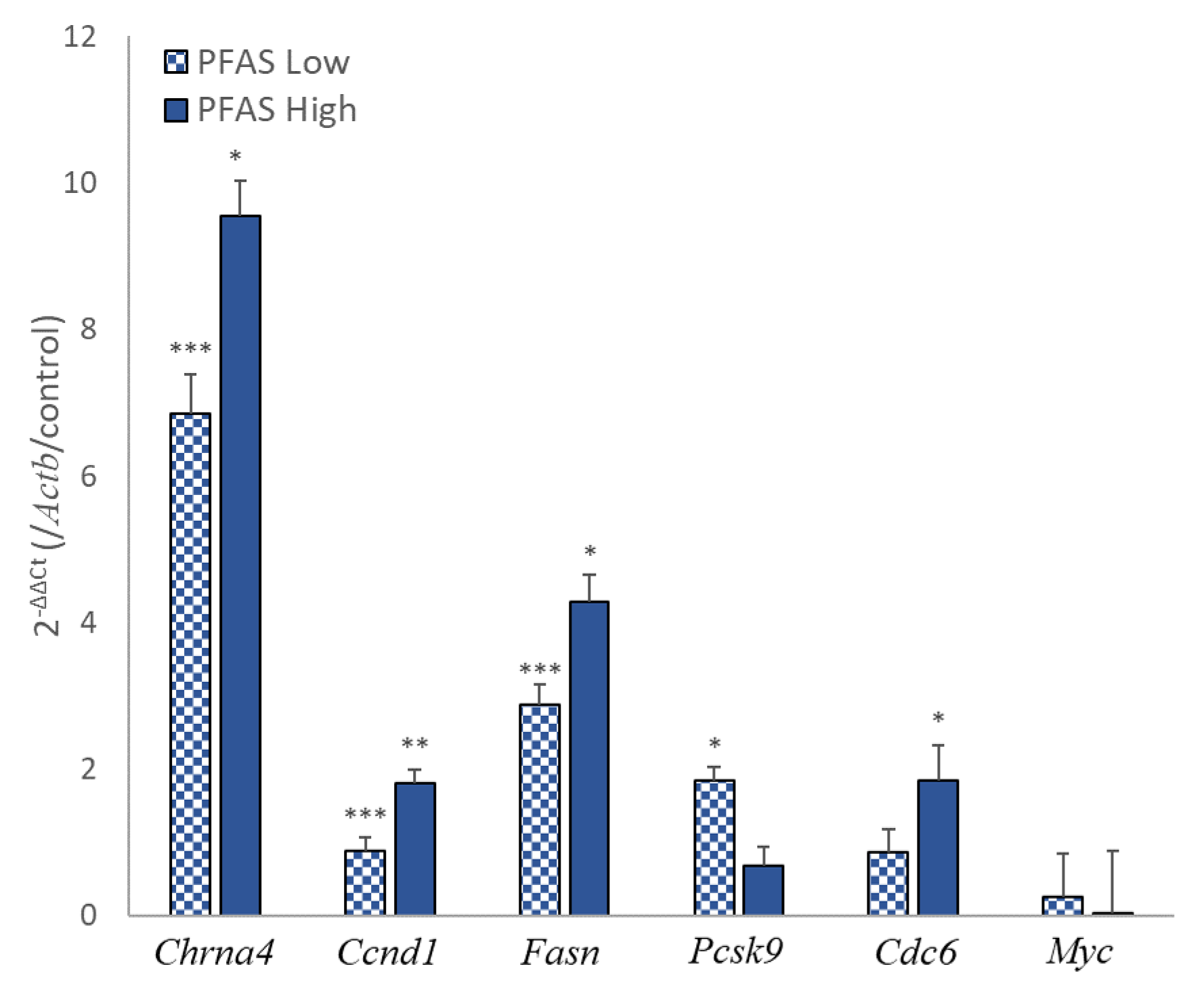
Real-time quantitative polymerase chain reaction. Representative genes of pathways with high variability in RNA sequencing were quantified. The y-axis shows the gene expression value relative to the control using β-actin (*Actb*) as the calibration gene. The data are represented as mean ± SEM (n = 5). All data were measured in duplicate. Asterisks (*) represent significant differences compared with control by Tukey–Kramer’s HSD test, *; *p* < 0.05, **; *p* < 0.01, ***; *p* < 0.001.

### 3.4. Metabolomics

Metabolome analysis was performed because it is a sensitive biomarker detection method for assessing the toxic effects of chemicals. In this analysis, 189 compounds were detected and quantified. The heat map of all samples and the cluster analysis based on the heatmap is shown in Fig. 4A. The control and PFAS-exposed groups were found to bifurcate at the beginning, and the metabolite variation due to PFAS exposure was similar in the PFAS-Low and PFAS-High groups. Enrichment analysis was performed for 121 of the 189 detected compounds listed in MetaboAnalyst R. Comparison of the PFAS-High and control groups revealed that the ether lipid pathway was the most significantly altered pathway, followed by the glycerophospholipid pathway. Comparison of the PFAS-Low and control groups showed different results, with the nucleotide glucose metabolism pathway being the most significantly altered pathway, followed by the glutathione metabolism pathway (Fig. 4B). The glycerophospholipid pathway is a synthetic pathway for the major components of cell membranes, whereas the ether lipid pathway involves a group of lipids derived from the glycerophospholipid pathway and has more physiological activity. The nucleotide sugar pathway is a glycosyl and phosphoglycosyl donor in the biosynthesis of carbohydrates and glycoconjugates in all living organisms. However, none of the compounds showed significant differences when their concentrations were compared between each group (FDR adjusted *p* > 0.05).

**Fig. 4.**
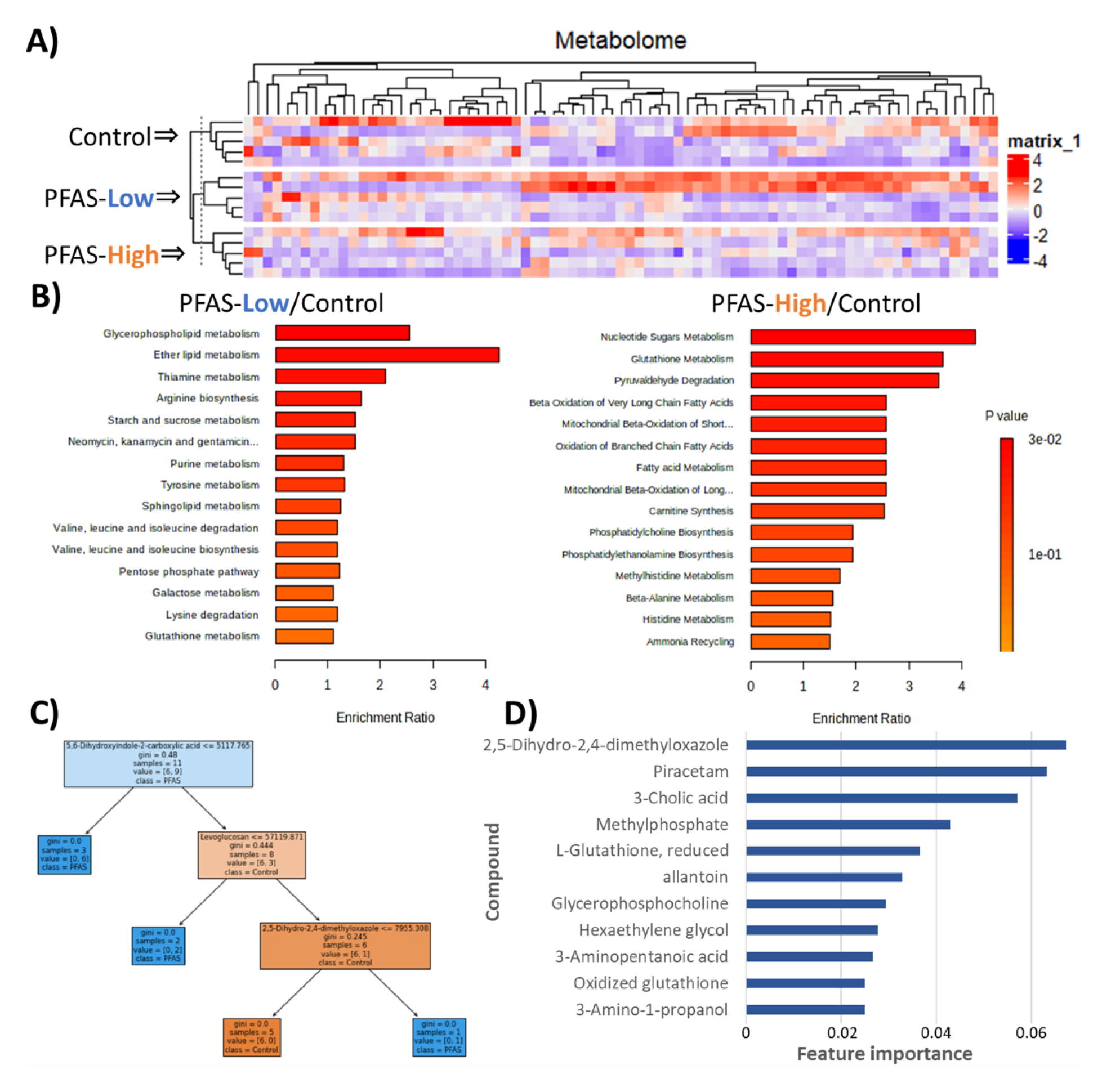
Metabolomics of the liver. A) Heatmap of all 189 metabolites detected in the liquid chromatography-mass spectrometry analysis and hierarchical clustering. B) Quantitative enrichment analysis compared with PFAS-High or PFAS-Low and control groups. C) An example of a decision tree in a random forest classifier. D) Top 10 feature importance for the random forest classifier (11 compounds were tied for the 10th place).

Hence, to characterize the effects of PFAS exposure on the liver metabolome, a binary classification of PFAS-exposed and control groups based on the metabolome using random forests was performed. The random forest classifier is a machine learning algorithm that constructs multiple decision trees at training time and outputs the class that is the mode of the classification or mean prediction of the individual trees. The method is suitable for classification using many features and is also appropriate for this research application as it allows visualization of the importance of each feature in the classification. The random forest classifier consisting of the decision tree shown in Fig. 4C discriminated PFAS-exposed and control groups from the metabolome with 100% accuracy for the training set and 90% accuracy for the test set. The importance of each feature (compound) in the classifier was calculated; 114 of the 189 compounds had a feature importance of 0, which means that they were not used in the classifier. The top 10 most important features are shown in Fig. 4D. The most important compound was 2,5-dihydro-2,4-dimethyloxazole. It is classified in the class of oxazolines and localizes to the cytoplasm, but there are almost no reports on its biological functions (Human Metabolome Database, 2021). Piracetam, the second most important substance, has been detected in the urine metabolome of patients with cancer, although its biological function as a metabolome remains unclear (Simón-Manso et al., 2019). The concentration profiles of these top 10 substances are shown in Fig. S2.

### 3.5. Molecular docking

The binding energy of each PFAS to PPAR, the receptor responsible for chemical-responsive hepatocyte hypertrophy, was evaluated by molecular docking. Known ligands for each PPAR were also docked for comparison (Jones et al., 2017; Oliveira et al., 2007; Tang et al., 2014) (Fig. 5, Table S2). PFOS (C8) exhibited the highest binding energy to all PPARα, PPARγ, and PPARδ, followed by PFNA (C9), PFOA (C8). Compared with known ligands for PPARs, compounds with carbon chain lengths above PFHxA (C6) showed higher docking scores with those known ligands and showed docking scores equal to or higher than those of the known ligands for PPARs. The linear correlation between these PFAS carbon chain lengths and docking scores showed a positive correlation with high linearity with R^2^ values of ≥0.80 (Fig. 5B). Comparing the binding poses of PFOA and known ligands of PPARs, PFOA was docked into the same binding pocket as the known ligands in all three PPARs (Fig. 5B). The number of amino acids in PPARs interacting with the ligands was fewer for PFOA than for the known ligands. However, the interacting amino acid species were similar. The binding poses of eight compounds other than PFOA are listed in Fig. S3.

**Fig. 5.**
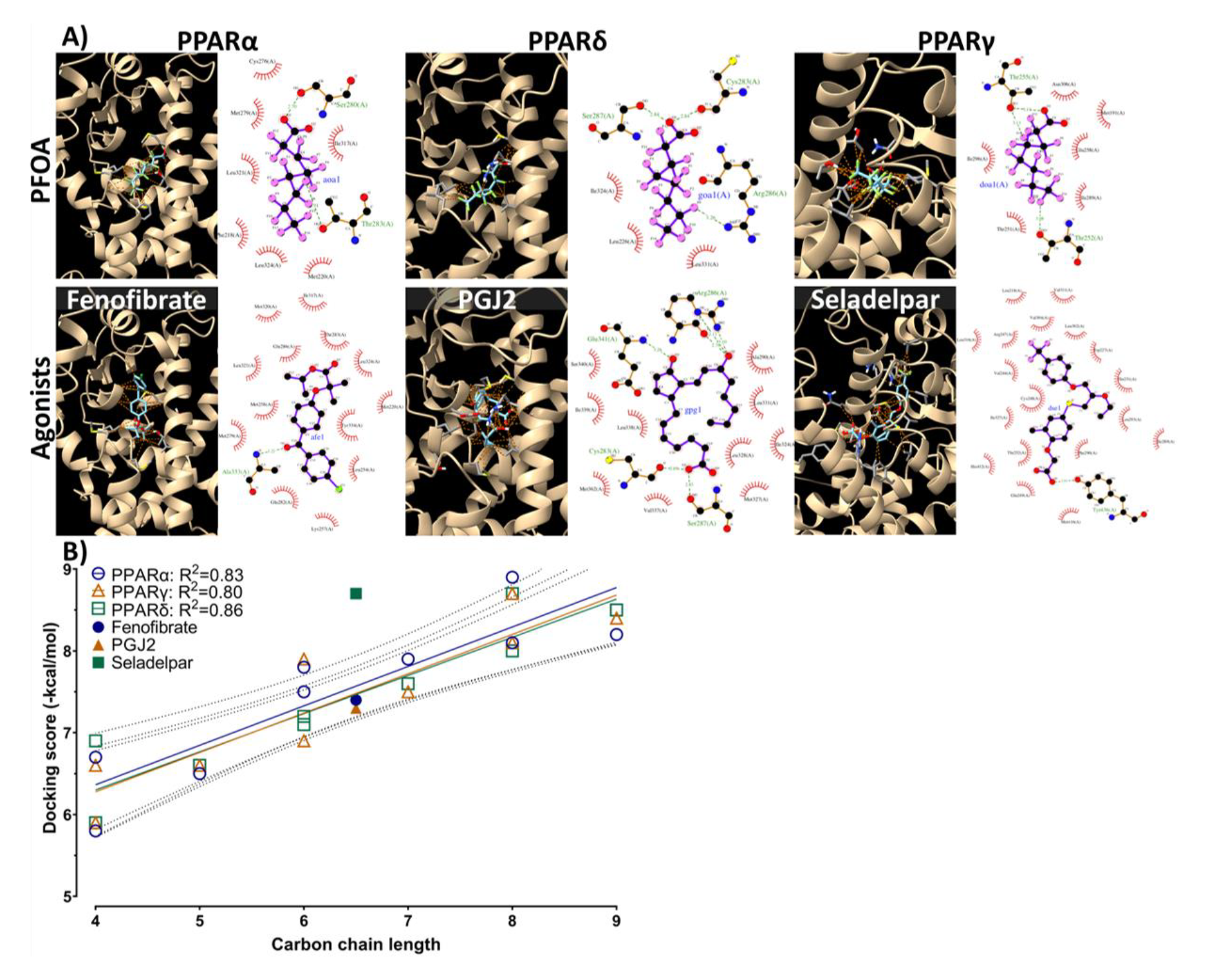
A) Theoretical binding pose obtained by molecular docking simulation for peroxisome proliferator-activated receptor (PPAR) with perfluorooctanoic acid or their known ligands. The protein is shown in ribbon representation with the binding residues shown in stick representation, and orange dashed lines in the 3D diagram indicate ligand–protein interactions. These interactions are visualized as 2D figures using LigPlot+. B) Regression of docking scores and carbon chain length of PFAS. The known ligands are indicated by filled shapes. Linear regression was performed using GraphPad Prism 9. Dashed lines represent the 95% confidence interval of the regression shown as colored lines. The docking scores are shown in Table S2.

### 3.6. Biomarkers for liver injury

To evaluate whether the PFAS exposure caused liver damage, levels of ALP and AST (common liver biomarkers) and cholesterol (a lipid metabolism marker) were measured using mouse plasma. Although all parameters were within the reference range in all three groups (Otto et al., 2016; Quimby and Luong, 2006), ALP level was significantly higher in the PFAS-High group than in the PFAS-Low group. On the other hand, cholesterol level was significantly lower in the PFAS-High group than in the control group.

## 4. Discussion

PFAS are ubiquitous within the human population, raising concerns regarding their impact on human health. The concentrations of PFOA and PFOS in the human plasma are known to range 1–30 ng/mL (Hölzer et al., 2021). It is imperative to undertake a comprehensive risk evaluation of actual low-concentration mixed exposures occurring within the real-world environment. Herein, mice were orally exposed to 1 µg/L (adequate to 100 ng/kg/day) of nine linear PFAS, including 25 ng/kg/day PFOA and 15 ng/kg/day PFOS, via drinking water for 5 weeks.

Histological examination revealed significant alterations in the liver (Figs. 1A and S1). These alterations were observed in both PFAS-Low (100 ng/kg/day) and PFAS-High (5,000 ng/kg/day) groups and were characterized by vacuolar degeneration featuring eosinophilic granules and anomalous arrangement of hepatic sinusoids. Additionally, hepatocyte vacuolation was frequently observed in the hepatic lobule, especially near the hepatic portal vein. This suggests that PFAS is taken up via the gastrointestinal tract and directly affects hepatocytes proximal to the portal vein. The gross abnormalities could be attributed to glycogen denaturation or chemical-responsive acidophilic granular hepatic hypertrophy. Given the findings of negative glycogen denaturation in PAS staining (Fig. 1B), chemical-responsive hepatic hypertrophy was presumed to be the etiology. Chen et al. (2022) postulated that inflammation and apoptosis of hepatocytes were demonstrable in response to PFOS exposure via drinking water at 500 µg/L, with Elcombe et al. (2012) positing that periportal hepatocellular vacuolation in rats could result from PFOS intake of 20 or 100 ppm. However, this study substantiated that even much lower mixed exposure doses could cause analogous histological alterations.

Omics analysis was performed to evaluate molecular biological alterations in the liver. Transcriptome analysis revealed a similar pattern of genetic variation between the PFAS-Low and -High groups (Fig. 2B). The genes were categorized according to their function based on the KEGG pathway, and the genes related to metabolism and carcinogenesis were found to be predominantly modified by PFAS exposure (Fig. 2C and D). Quantitative PCR was performed to validate the changes in the expression levels within these pathways. *Chrna4*, the most highly expressed protein, was upregulated approximately 10-fold in the high-exposure group and 6-fold in the low-exposure group compared with the untreated control group (Fig. 3). *Chrna4* is located upstream of *Ccnd1*, an oncogene in the KEGG chemical carcinogenesis pathway, suggesting that *Chrna4* contributes to chemical carcinogenesis. *Fasn*, which encodes a fatty acid synthase, was also significantly upregulated. The overexpression of *Fasn* and enhanced lipid metabolism promote cancer cell growth and metastasis as cancer cells utilize fatty acids for tumor growth and metastasis (Fhu and Ali, 2020). These results suggest that PFAS exposure enhances lipid metabolism and promotes tumorigenesis. *Ccnd1* and *Cdc6* are regulators of cell proliferation in the cell cycle. *Ccnd1* triggers the G1–S phase transition in the cell cycle by activating cyclin-dependent kinases (*Cdk4* and *Cdk6*) and enhanced cancer cell proliferation (Valla et al., 2022). *Cdc6*, a cell growth regulator, is expressed in the quiescent phase (G1 phase) in normal cells but is expressed at all cell cycle stages in cancer cells and enhances cancer cell proliferation. The increased expression of these genes in this study indicates that even low concentrations of PFAS mixture exposure might increase the risk of carcinogenesis.

Metabolome analysis revealed similar patterns of metabolites altered by PFAS in the PFAS-Low and -High groups (Fig. 4A). On the other hand, enrichment analysis suggested that different pathways were altered in these groups, with the glycerophospholipid and ether lipid pathways being particularly affected in the PFAS-Low group. The main factor contributing to this is the decrease in glycerophosphocholine (α-GPC) levels in the PFAS-Low group (Fig. S4A). This suggests that following PFAS exposure, the decrease in acetylcholine precursor α-GPC levels may have led to an increase in the expression of the acetylcholine receptor Chrna4 (Fig. 3) to compensate for the initial decrease. The glutathione metabolic pathway, a defense mechanism against oxidative stress, was upregulated in the PFAS-High group. The nucleotide sugar pathway was upregulated in the both PFAS-exposed group owing to an increase in glucose 6-phosphate levels (Fig. S4B); G6P is the starting substrate for the pentose phosphate pathway, a NADPH-producing pathway in the liver. The increase may have occurred in response to the NADPH requirement of glutathione. On the other hand, the increase in reduced and oxidized glutathione levels (Fig. S2) suggests that the oxidative stress response was not disrupted and that the glutathione level increased adaptively. It should be noted that not all of the individual variations for each of these compounds were statistically significant. While the enrichment analysis showed different results among the exposure groups, the binary classification by machine learning, in addition to cluster analysis, could discriminate PFAS exposure from the metabolome with 90% accuracy, suggesting that there are exposure-specific effects. The substances that contributed most to the classification, 2,5-dihydro-2,4-dimethyloxazole and piracetam, have unknown physiological significance. Further analysis of these substances may reveal the complete picture of the AOP of PFAS. The decision tree machine learning method to classify exposure groups effectively evaluated toxicity effects in a *p*-value-independent manner based on compounds with unknown biological functions. On the other hand, the ether lipid pathway obtained by RNA-seq analysis did not show significant changes in gene expression (data not shown). These findings suggest that multi-omics analysis may be advantageous in detecting prepathological changes.

One of the etiological factors underlying hepatic precancerous proliferative alterations is the activation of nuclear receptors, PPARs. In mammals, three types of PPARs have been identified, PPARα, PPARγ, and PPARδ, which participate in cell differentiation, energy metabolism, and cancer cell growth (Cheng et al., 2021). Molecular docking simulations were performed to evaluate the binding affinity between nine PFAS and mouse PPARα, PPARγ, and PPARδ. The results indicated a robust correlation between the length of the PFAS carbon chain and binding affinity for all three PPAR isotypes in mice (Fig. 5B). Notably, PFOS, PFNA, and PFOA had binding poses comparable to that of fenofibrate, a known PPARα ligand, and exhibited high binding potency exceeding that of fenofibrate (Table S2), suggesting that PFAS are PPARα agonists. Conversely, short-chain PFAS could also be bioactive, as docking scores equal to or above −7.0 generally exhibit high pharmacological activity in nanomolar order in drug discovery screening (Kuenemann et al., 2018). A previous study reported that PFOA, PFOS, and short-chain PFAS could activate PPARα and PPARγ *in vitro* (Evans et al., 2022). Short-chain PFAS have relatively low toxicity but may contribute to PPAR-dependent hepatic hypertrophy. Although hepatocyte hypertrophy was observed, the RNA-seq results indicated that the expression levels of PPAR-responsive genes were not remarkably elevated (e.g., 2.0-fold increase in *Cyp7a1* expression in the PFAS-High groups). However, an increased expression of cell proliferative genes, including *Ccnd1*, in the oncogenic pathway indicated the presence of other receptor-binding-mediated cell proliferation signals. To accurately assess the toxicity of PFAS, key AOP events related to the hepatocyte hypertrophy require elucidation.

In establishing regulatory limits, it is crucial to discern the indications of toxic effects. Therefore, plasma ALP, AST, and cholesterol levels were measured as biomarkers of liver injury (Fig. 6). Although ALP levels increased significantly following exposure to high concentrations of PFAS, they remained within the normal range. AST and cholesterol levels were within the normal range and exhibited a decreasing trend correlated with increasing exposure. These findings suggest that PFAS exposure did not cause severe liver injury, even though histological alterations in the liver were evident. Hepatocytes transiently enlarge as an adaptive response to chemical exposure, whether adverse or nonadverse (Hall et al., 2012); therefore, the histological changes are considered adaptive hypertrophy. Because carcinogenic genetic changes were observed in this study, there remains a risk of liver damage with prolonged exposure. Therefore, chronic toxicity studies of mixed exposures are essential to elucidate the point.

**Fig. 6.**
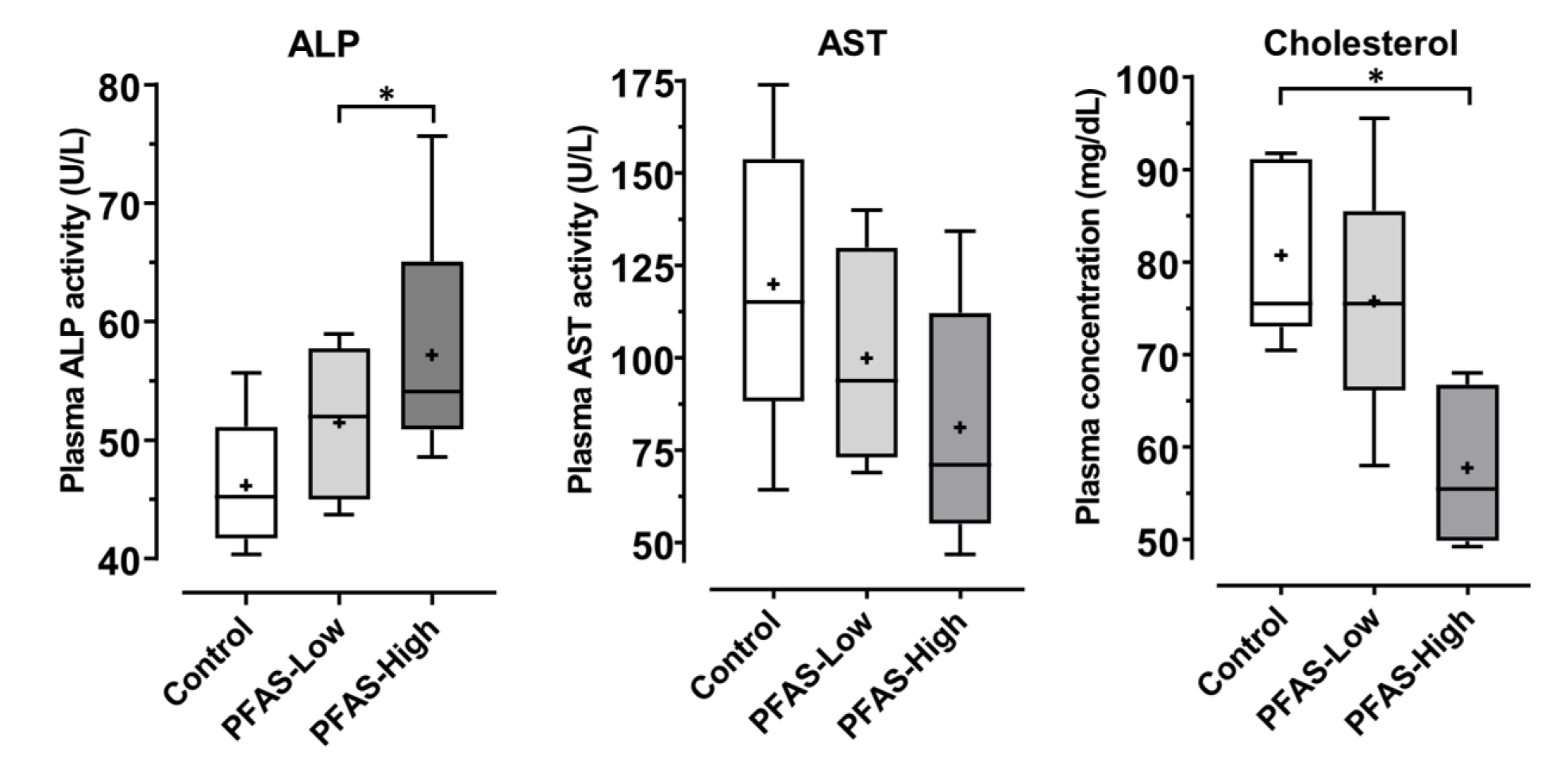
Plasma concentration of hepatic biomarkers. The plasma concentrations of ALP (alkaline phosphatase), AST (aspartate aminotransferase), and cholesterol in plasma were measured colorimetrically. Data are presented in box-and-whisker plots, with boxes indicating each quartile and median, + denoting the mean value, and whiskers denoting the maximum and minimum values. All data were measured in duplicate. * represents significant differences detected by Tukey–Kramer’s HSD test (*p* < 0.05).

Although transient changes improve as PFAS are excreted, they are considered highly hazardous owing to their long half-lives. Moreover, species differences should be considered when extrapolating these findings to humans. This study utilized mice as test animals. A previous study showed that the sensitivity of PPARα to PFAS is higher in mice than in humans and that PFAS at levels that cause significant carcinogenicity in rodents may be insensitive or unresponsive in humans (Cheung et al., 2004). However, the half-lives of PFOA, PFOS, and other PFAS compounds are shorter in mice than in humans, and the same exposure may result in higher concentrations of residues in humans (Fenton et al., 2021). Therefore, it is essential to consider more appropriate models to evaluate the health effects of PFAS in humans. One limitation of this study is that it involved mixed exposure; therefore, it is unknown whether there were individual effects of each substance or synergistic effects. Short-chain PFAS are alternatives to PFOA and PFOS because of their shorter half-lives. Their lower binding to each PPAR than long-chain PFAS suggest that they are less toxic. However, previous studies have shown that PFBS, a short-chain PFAS, can be toxic. For instance, a single administration of 300 mg/kg PFBS to mice caused PPAR-responsive hepatic gene alteration (Lau et al., 2020). Drinking water exposure to 10 µg/L PFBS significantly decreased phospholipid levels in the murine liver (Chen et al., 2022). Therefore, short-chain PFAS can result in toxicity (like PFOA and PFOS) upon persistent exposure.

## 5. Conclusions

This study confirmed that low concentrations of PFAS mixtures can cause changes in gene expression (including cancer-related genes) and histology, even after a relatively short exposure period of 5 weeks. Thus, it is essential to conduct exposure studies for longer periods and at lower concentrations that reflect environmental exposure conditions.

## Abbreviations^1^

^1^AOP: Adverse outcome pathway
CI: Confidence interval
DNEL: Derived no-effect level
EFSA: European Food Safety Authority
EPA: Environmental Protection Agency
FC: Fold change
FDR: False discovery rate
HE: Hematoxylin and eosin
KEGG: Kyoto Encyclopedia of Genes and Genomes
MHLW: Ministry of Health, Labour and Welfare
PAS: Periodic acid-Schiff
PFAS: Poly-fluoroalkyl substances
PFBA: Perfluorobutanoic acid
PFBS: Perfluorobutanesulfonic acid
PFHpA: Perfluoroheptanoic acid
PFHxA: Perfluorohexanoic acid
PFHxS: Perfluorohexanesulfonic acid
PFNA: Perfluorononanoic acid
PFOA: Perfluorooctanoic acid
PFOS: Perfluorooctanesulfonic acid
PFPeA: Perfluorovaleric acid
PPAR: Peroxisome proliferator-activated receptor
PCR: Polymerase chain reaction
PXR: Pregnane X receptor
QC: Quality control
QTOF-MS: Quantitative time-of-flight mass spectrometry
UPLC: Ultra-performance liquid chromatography

## Acknowledgments

This work was supported by Grants-in-Aid for Scientific Research from the Ministry of Education, Culture, Sports, Science and Technology of Japan awarded to M. Ishizuka (Nos. 22KK0163 and 21H04919) and K. Takeda (Nos. 23K14091, 21K14988, and 20J01553). This work was partly supported by JST OPERA Program Japan and an unrestricted grant of Yamada Bee Company, Japan awarded to C. Mori for the metabolome analysis. We would also like to thank Enago (Crimson Interactive Pvt. Ltd. Mumbai, Maharashtra, Republic of India) for editing and proofreading this manuscript.

