## Supplemental material for "Subchronic Toxicity Assessment of Perfluoroalkyl Substances by Mixed Exposure of Nine Substances at Daily Intake Relevant Concentration"

**Supplementary material**

Fig. S1. Histological examination of the kidney, heart, and spleen by hematoxylin and eosin staining.

Fig. S2. Box-and-whisker plot of the top 10 essential compounds in the random forest classifier.

Fig. S3. Representative 2D diagram of binding poses between peroxisome proliferator-activated receptors (α, δ, and γ) and poly-fluoroalkyl substances (PFOS, PFHxS, and PFBS) obtained by molecular docking.

Fig. S4. Box-and-whisker plot of glycerophosphocholine and D-glucose 6-phosphate, which contributed to the enrichment analysis shown in Fig. 5B.

Table S1. Primer information

Table S2. Docking scores

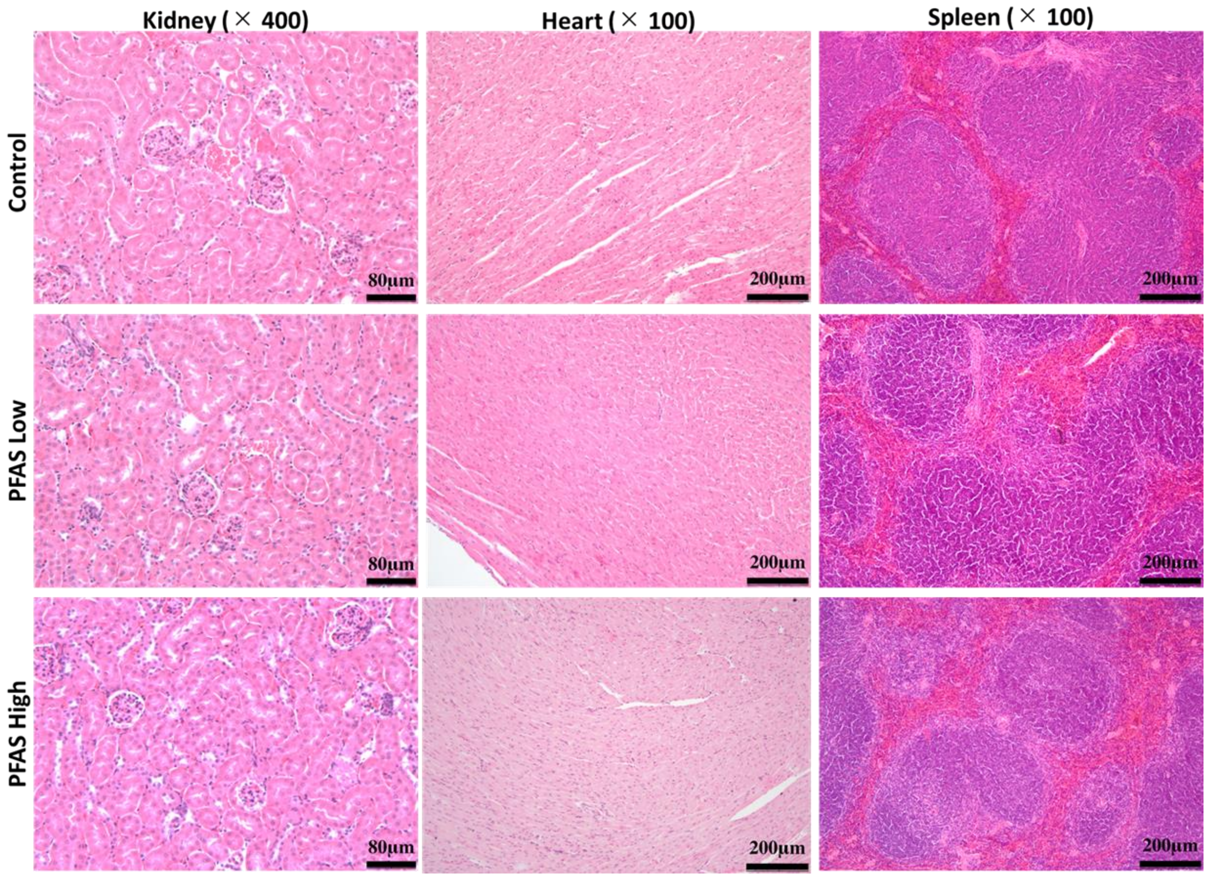


**Fig. S1.** Histological examination of the kidney, heart, and spleen by hematoxylin and eosin (HE) staining.

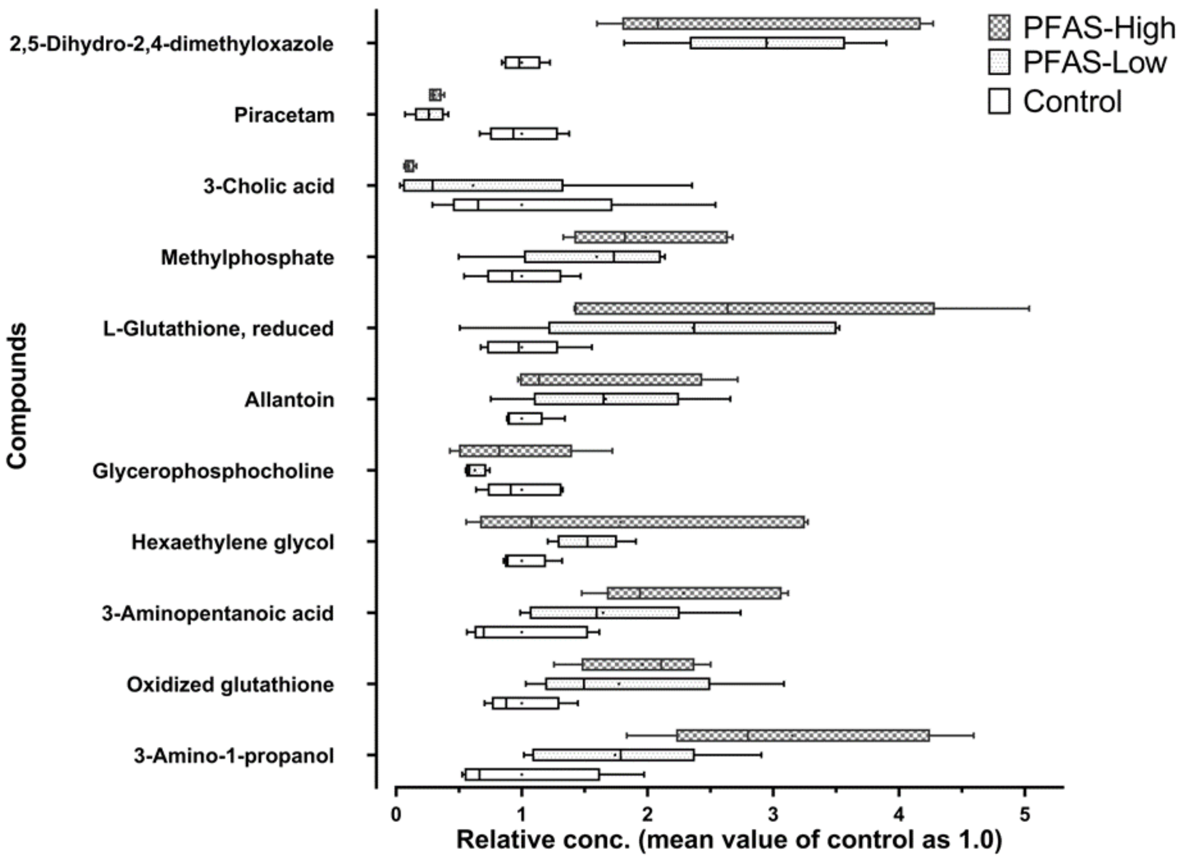


**Fig. S2.** Box-and-whisker plot of the top 10 essential compounds in the random forest classifier shown in Fig. 4D. The x-axis represents the relative concentration corrected by dividing it by the control group's mean for each compound (the mean of the control should be 1.0). + indicates mean values. No significant differences were detected in all substances (FDR adjusted $p > 0.05$).

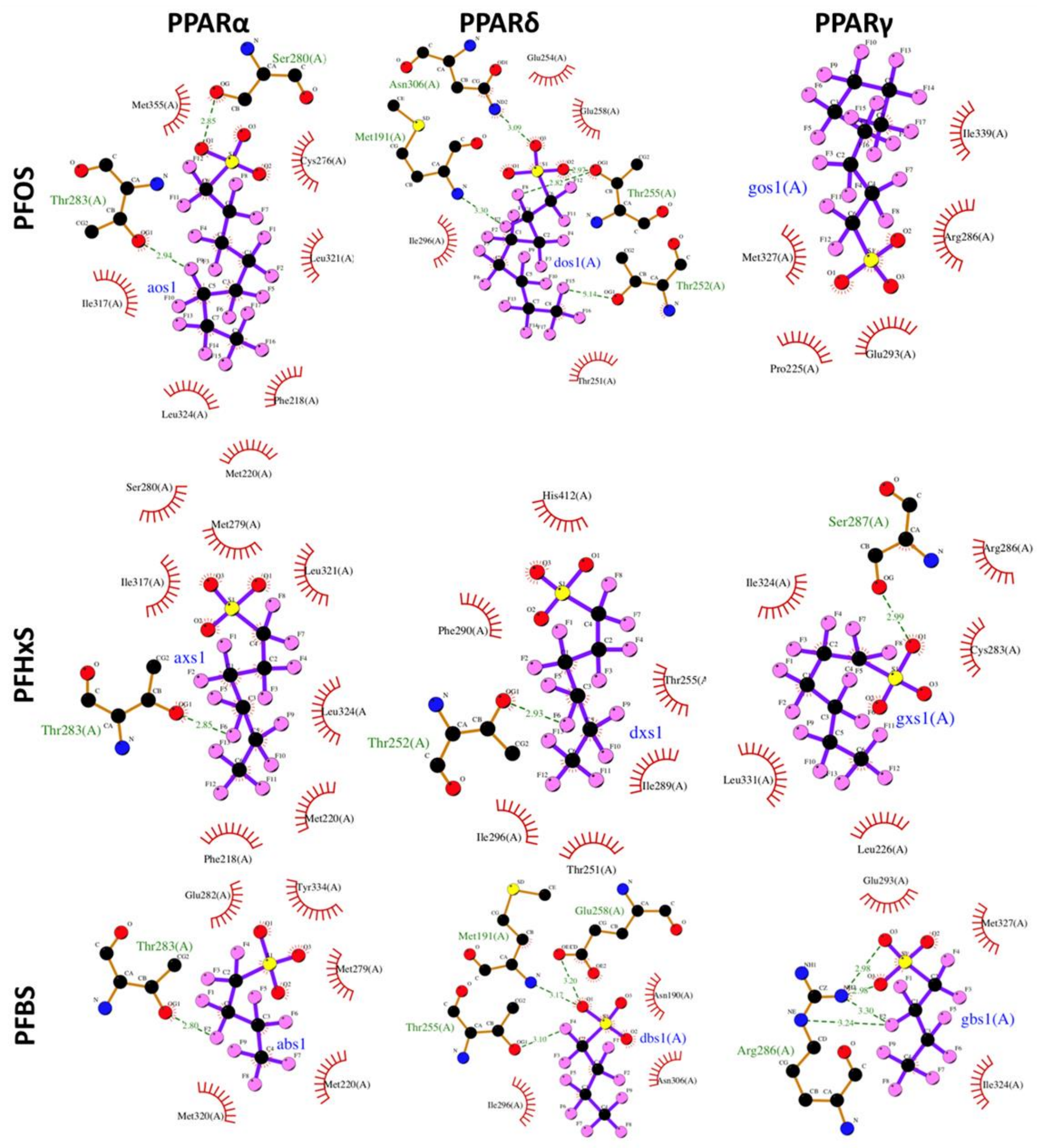


**Fig. S3.** Representative 2D diagram of binding poses between peroxisome proliferator-activated receptors (α, δ, and γ) and poly-fluoroalkyl substances (PFOS; perfluorooctanesulfonic acid, PFHxS; Perfluorohexanesulfonic acid, and PFBS; perfluorobutanesulfonic acid) obtained by molecular docking.

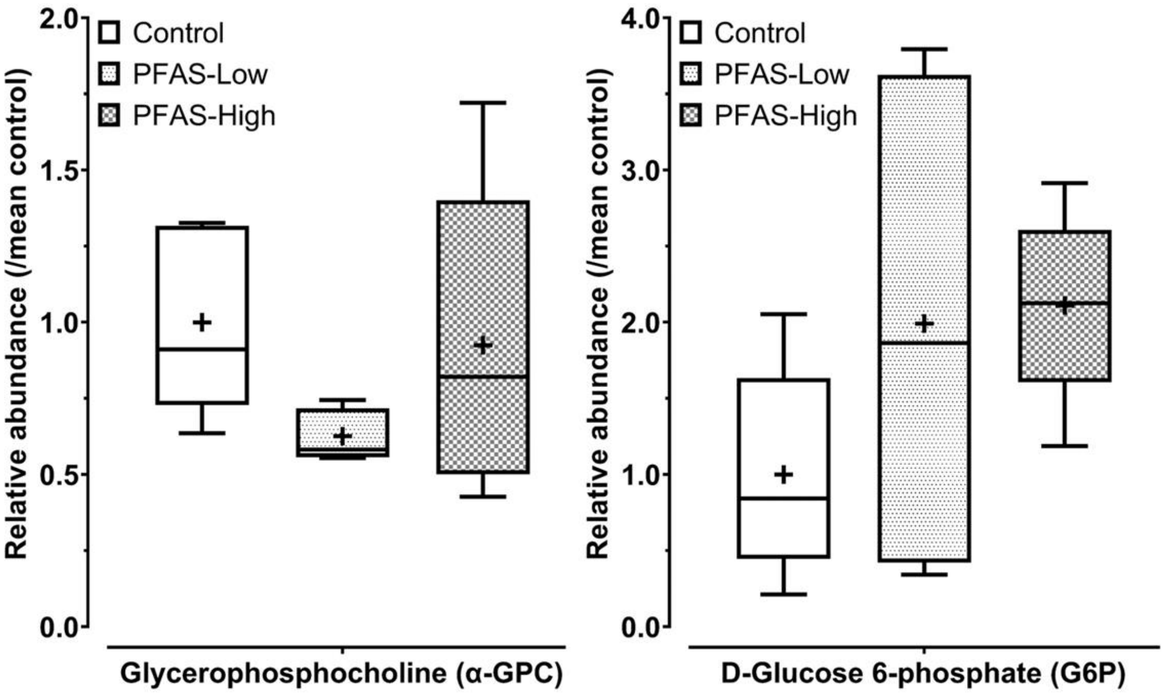


**Fig. S4.** Box-and-whisker plot of glycerophosphocholine (α-GPC) and D-glucose 6-phosphate, which contributed to the enrichment analysis shown in Fig. 4B. The y-axis represents the relative concentration corrected by dividing it by the mean of the control group. + indicates mean values. No significant differences were detected in all substances (false discovery rate adjusted $p > 0.05$).

**Table S1. Primer information**

| Gene (Product size) | Sequences | Amplification efficiency (%) Standard error of mean |
|---|---|---|
| ***Chrna4*** | Forward：GTGTGGGTGAAGCAGGAGTG | 103.4 |
| (177 bp) | Reverse：GGTGGGTGACTGCAAAGTCC | 3.2 |
| ***Ccnd1*** | Forward：CATTTCCAACCCACCCTCCA | 97.9 |
| (152 bp) | Reverse：CCAGGGCCTTGACCGGG | 4.5 |
| ***Pcsk9*** | Forward：ATCACCGACTTCAACAGCGT | 90.2 |
| (174 bp) | Reverse：GCCCTTCCCTTGACAGTTGA | 2.3 |
| ***Fasn*** | Forward：CCTTCGGTTCAGTCTCTTTCCA | 90.6 |
| (111 bp) | Reverse：ACACCCTCCAAGGAGTCTCAC | 1.7 |
| ***Cdc6*** | Forward：TTACGGTGGTGATCGAGACG | 96.9 |
| (103 bp) | Reverse：TGATGGCCACACAACTCTCTG | 4.3 |
| ***Myc*** | Forward：GTTGGAAACCCCGCAGACAG | 99.1 |
| (94 bp) | Reverse：ATAGGGCTGTACGGAGTCGT | 3.5 |
| ***Actb*** | Forward：ACTGTCGAGTCGCGTCCA | 97.9 |
| (86 bp) | Reverse：ATCCATGGCGAACTGGTGG | 0.6 |

All primers were designed using the National Center for Biotechnology Information Primer designing tools. Amplification efficiency was determined by six points of 5-fold step dilution (number of replicates = 4).

**Table S2.** Docking scores

| Compound | C length | Docking score (kcal/mol) | | |
|---|---|---|---|---|
| | | PPARα | PPARγ | PPARδ |
| PFOS | C8 | 8.9 | 8.7 | 8.7 |
| PFNA | C9 | 8.2 | 8.4 | 8.5 |
| PFOA | C8 | 8.1 | 8.1 | 8 |
| PFHpA | C7 | 7.9 | 7.5 | 7.6 |
| PFHxS | C6 | 7.8 | 7.9 | 7.2 |
| PFHxA | C6 | 7.5 | 6.9 | 7.1 |
| PFBS | C4 | 6.7 | 6.6 | 6.9 |
| PFPeA | C5 | 6.5 | 6.6 | 6.6 |
| PFBA | C4 | 5.8 | 5.9 | 5.9 |
| Agonists | - | 7.4 | 7.3 | 8.7 |

Theoretical docking scores were calculated by molecular docking. The representative docking poses are shown in Fig. 5 and S3. Typical agonists of each peroxisome proliferator-activated receptor (PPAR) were docked into the PPARs for comparison. Agonists for PPARα; fenofibrate, PPARγ; prostaglandin J2, PPARδ; seladelpar.
